# How stimulation waveform shape affects collective oscillations in the brain networks

**DOI:** 10.64898/2026.06.16.732561

**Authors:** Vivek Sharma, Paul H. E. Tiesinga, Joana Cabral

## Abstract

Brain oscillations emerge from nonlinear interactions across anatomically connected neural populations, alternating between transiently coordinated and desynchronised states. Transcranial alternating current stimulation (tACS) can modulate these dynamics, but most work has focused on frequency and amplitude, leaving waveform shape comparatively unexplored. Here we used a whole-brain model of delay-coupled Stuart–Landau oscillators constrained by empirical human structural connectivity to determine how sinusoidal, square, triangular, sawtooth and pulsed stimulation reshape spontaneous alpha-band activity. All waveforms were applied at the same frequency and amplitude to the posterior parieto-occipital regions. Network responses were quantified using the Kuramoto order parameter, spectral entropy and metastable oscillatory modes of transient alpha bursts. Sinusoidal and pulsed stimulation produced the strongest effects, increasing global synchrony while reducing metastability and spectral entropy, consistent with a transition from a fluctuation-rich regime to a coherent and spectrally concentrated state. These waveforms also transformed intermittent alpha bursts into prolonged or near-continuous oscillatory episodes. In contrast, square, triangular and sawtooth stimulation reduced synchrony while largely preserving metastability, producing weaker and more fragmented modulation. These findings identify waveform shape as a key determinant of rhythmic stimulation effects and a principled parameter for neuromodulation design.

## Introduction

Endogenous brain rhythms play a fundamental role in coordinating perception, cognition, and behaviour across spatial and temporal scales [1, 2]. Non-invasive brain stimulation techniques such as transcranial alternating current stimulation (tACS) provide a powerful means to probe and modulate these oscillations by applying weak, rhythmic electric fields to the cortex [3]. Traditionally, tACS has been delivered as a sinusoidal current, entraining neural populations at frequency-specific rhythms (e.g., in the *α* band) and thereby offering both mechanistic insight into brain function and potential clinical applications in disorders of dysrhythmia [4, 5].

Recent advances in stimulation hardware have expanded the available parameter space, enabling the delivery of arbitrary waveforms beyond the canonical sinewave. This development has prompted growing interest in how waveform shapes, including square, triangular, sawtooth, and pulsed currents, may differentially modulate neural activity. Theoretical and experimental evidence indicate that waveform shape influences entrainment efficacy: sharper slopes or discontinuities provide stronger depolarising drives, increasing the probability of neuronal firing compared to smoothly varying sinusoids [6, 7]. For instance, positive-ramp sawtooth stimulation at *α* frequency has been shown to more effectively enhance ongoing oscillations than either sinusoidal or negative-ramp variants [6]. Similarly, triangular or pulsed waveforms have been reported to reorganise network connectivity or enhance corticospinal excitability in ways that differ from sinusoidal tACS [8]. These findings suggest that waveform shape is not a trivial parameter but a key determinant of how external rhythms interact with endogenous oscillatory dynamics.

Despite these advances, the systematic effects of waveform shape on large-scale brain network dynamics remain poorly understood. Human EEG/MEG studies face challenges in disentangling stimulation artifacts from neural signals, while empirical findings on waveform-specific effects are often mixed and limited to local or task-specific measures. Computational whole-brain models, which integrate structural connectivity with local oscillatory dynamics, provide a complementary framework for testing how perturbations of different forms propagate across the connectome and shape emergent network activity [9, 10]. Such models allow for controlled comparisons of sinusoidal and non-sinusoidal stimulation, yielding mechanistic predictions that can inform experimental design and translational applications.

With a dynamical-systems perspective in view, whole-brain activity can be viewed as the motion of a high-dimensional system on a landscape of metastable attractors shaped by structural coupling and local bifurcation parameters [11, 12]. Small, weakly structured inputs probe the system’s linear susceptibility (how readily phases align), whereas temporally structured perturbations can transiently reconfigure basins of attraction, revealing state-dependent critical slowing down and differences in global stability. In this view, entrainment by transcranial wave-forms reflects controlled phase-response interactions with local Hopf–like nodes embedded in a delayed, diffusively coupled network. In addition to input strength, waveform shape modulates not only resonance at the target frequency but also the recruitment of nonlinear harmonics and cross-scale interactions. Building on in-silico perturbation studies that distinguished brain states via recovery latencies and integration dynamics, this work tests whether the shape of periodic drive provides an independent handle on network susceptibility—altering alpha-band synchrony, spectral concentration, and metastability in ways that standard sinusoidal stimulation cannot [10, 13].

Here, we use a whole-brain dynamical model based on coupled Stuart–Landau oscillators to investigate how the shape of externally applied waveforms modulates network activity. By systematically comparing sinusoidal, square, triangular, pulsed, and sawtooth perturbations, we examine their impact on spectral power, phase synchrony, metastability, and spectral entropy, metrics that capture both local and global aspects of oscillatory brain dynamics. We hypothesise that waveform shape differentially modulates the dynamical regime of large-scale brain activity, such that distinct temporal profiles of stimulation can drive transitions between metastable and coherent oscillatory states even when frequency and amplitude are held constant. In doing so, we aim to provide a mechanistic account of how non-sinusoidal waveform tACS shapes brain dynamics, bridging theoretical models with experimental evidence and highlighting waveform shape as a critical parameter for future neuromodulation research.

## 1 Methods

### 1.1 Empirical data and functional connectivity

The empirical resting-state MEG data and anatomical connectivity analysed in this study were derived from Human Connectome Project (HCP; https://www.humanconnectome.org) and provided by Castaldo, Santos et al. (2023) [14]. For all preprocessing and data preparation steps, we refer the reader to [14]. The dataset comprises source-reconstructed resting-state MEG recordings from 89 healthy participants (6-minute eyes-open condition), together with population-averaged structural connectivity and inter-regional distance matrices derived from diffusion MRI.

Functional connectivity (FC) was quantified using amplitude-envelope correlation (AEC) [15, 16]. For each participant, the source-reconstructed regional time series from the 90-region Automated Anatomical Labeling (AAL90) atlas [17], comprising both cortical and subcortical regions, were band-pass filtered in the alpha band (8–13 Hz), and analytic signal envelopes were extracted via the Hilbert transform. Pairwise Pearson correlations between amplitude envelopes yielded participant-level AEC matrices, which were subsequently averaged across participants to obtain a group-level empirical FC matrix. This empirical FC served as the reference target for fitting the whole-brain model.

### 1.2 Whole-brain network model

Large-scale brain dynamics were modelled using a network of coupled oscillatory units, each representing a brain region. This approach builds on the theoretical framework in which local neural masses are described by a canonical Hopf normal form operating close to a bifurcation point [10, 14]. In this regime, each region exhibits noise-driven fluctuations that can give rise to damped or self-sustained oscillations depending on which side of the bifurcation it operates. Such a formulation captures the dual capacity of cortical assemblies to remain quiescent in the absence of stimulation while producing rhythmic activity when perturbed.

Formally, the dynamics of each node *n* are governed by

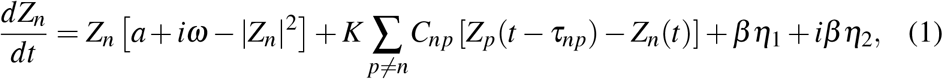

where *Z*_*n*_ is the complex state variable of node *n, a* controls the distance to the Hopf bifurcation, *ω* is the natural frequency, set to 40 Hz and *β* scales the stochastic drive. Nodes are coupled through the structural connectivity matrix *C*_*np*_, with conduction delays *τ*_*np*_ proportional to empirical fibre lengths. The global coupling constant *K* regulates the balance between local dynamics and long-range interactions.

Operating near the Hopf bifurcation ensures that the system is particularly sensitive to noise fluctuations, allowing the emergence of coherent envelope dynamics in specific frequency bands. This behaviour has been shown to reproduce key empirical features of resting-state MEG activity, such as frequency-specific amplitude correlations and metastable network organisation [10, 14]. By systematically varying the global coupling and conduction velocity parameters, we generated ensembles of simulations spanning the relevant dynamical regimes, from incoherence to strong synchronisation.

### 1.3 Model fitting

The whole-brain model was fitted to empirical data by comparing simulated functional connectivity (FC) with empirical FC matrices derived from amplitude envelope correlation (AEC). Fitting accuracy was quantified using the Pearson correlation coefficient between simulated and empirical FC, computed for each parameter combination of global coupling strength (*G*_*C*_) and mean conduction delay (*M*_*D*_).

Although the intrinsic frequency of each node was fixed at 40 Hz, slower collective fluctuations in the alpha band emerge naturally at the network level through delay-mediated interactions [9]. To ensure physiological relevance, both simulated and empirical signals were band-pass filtered in the alpha range (8–13 Hz), and FC was computed from the corresponding amplitude envelopes.

Across the parameter space, increasing global coupling and conduction delays led to a transition from desynchronised dynamics to the emergence of correlated power fluctuations in the alpha band, reflecting large-scale coordination mediated by time delays [9, 14]. Parameter sets yielding the highest correlation values within this band were identified as candidate fits.

The best-fitting parameter set achieved a correlation of *r* = 0.477 between simulated and empirical FC.

### 1.4 Driven Hopf Network

To investigate how different stimulation waveforms interact with large-scale brain dynamics, we introduced an external forcing term into the network equations. Perturbations were modelled as additive complex inputs to the derivative of the state variable, applied to a subset of nodes representing stimulated brain regions. Perturbations were applied to posterior visual-parietal regions associated with alpha-band dynamics [18, 19], corresponding anatomically to the bilateral cuneus, superior occipital cortex, superior parietal cortex, and precuneus (AAL regions; Figure 2).

Zaehle et al. (2010) demonstrated that alternating current stimulation applied bilaterally over parieto–occipital sites enhances endogenous alpha activity and entrains ongoing oscillations [18], indicating that these occipital areas are highly susceptible to rhythmic external drive. Motivated by these findings, we targeted the corresponding posterior visual–parietal nodes in the whole-brain model to examine how waveform shape modulates the entrainment of emergent alpha-band dynamics. The perturbed system is thus described as:

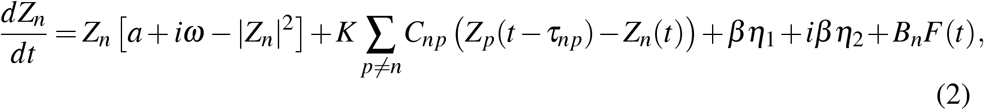

where *B*_*n*_ ∈ {0, 1} selects the stimulated nodes and *F*(*t*) denotes the waveform of the applied perturbation.

Here, *F*(*t*) defines the temporal profile of the stimulation waveform. The perturbation acts directly on both the real and imaginary components of the complex state variable, thereby modulating both the amplitude and phase of the local oscillation. Figure 3 shows an example of how a localised perturbation propagates through the network and alters the collective dynamics.

#### Perturbation waveforms

To systematically compare the influence of stimulation shape on whole-brain dynamics, we considered five canonical classes of periodic forcing: sinusoidal, square, triangular, pulsed, and sawtooth (Fig. 1). In all simulations, the external input was implemented as a complex-valued periodic forcing applied to the stimulated nodes,

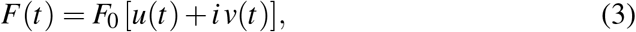

where *F*_0_ denotes the stimulation amplitude, *f* the stimulation frequency, and the real and imaginary components (*u, v*) define the waveform shape in quadrature. This formulation ensures that the perturbation acts on both the real and imaginary components of the local oscillator state. The equations below describe the waveforms used in the final simulations. Here, *t* is expressed in seconds, *f* is the stimulation frequency, *ϕ*(*t*) = frac( *ft*) is the phase within each cycle, and *ϕ*_*i*_(*t*) = frac( *ft* + 0.25) denotes the quadrature-shifted phase. The following waveform classes were examined:

**Figure 1.**
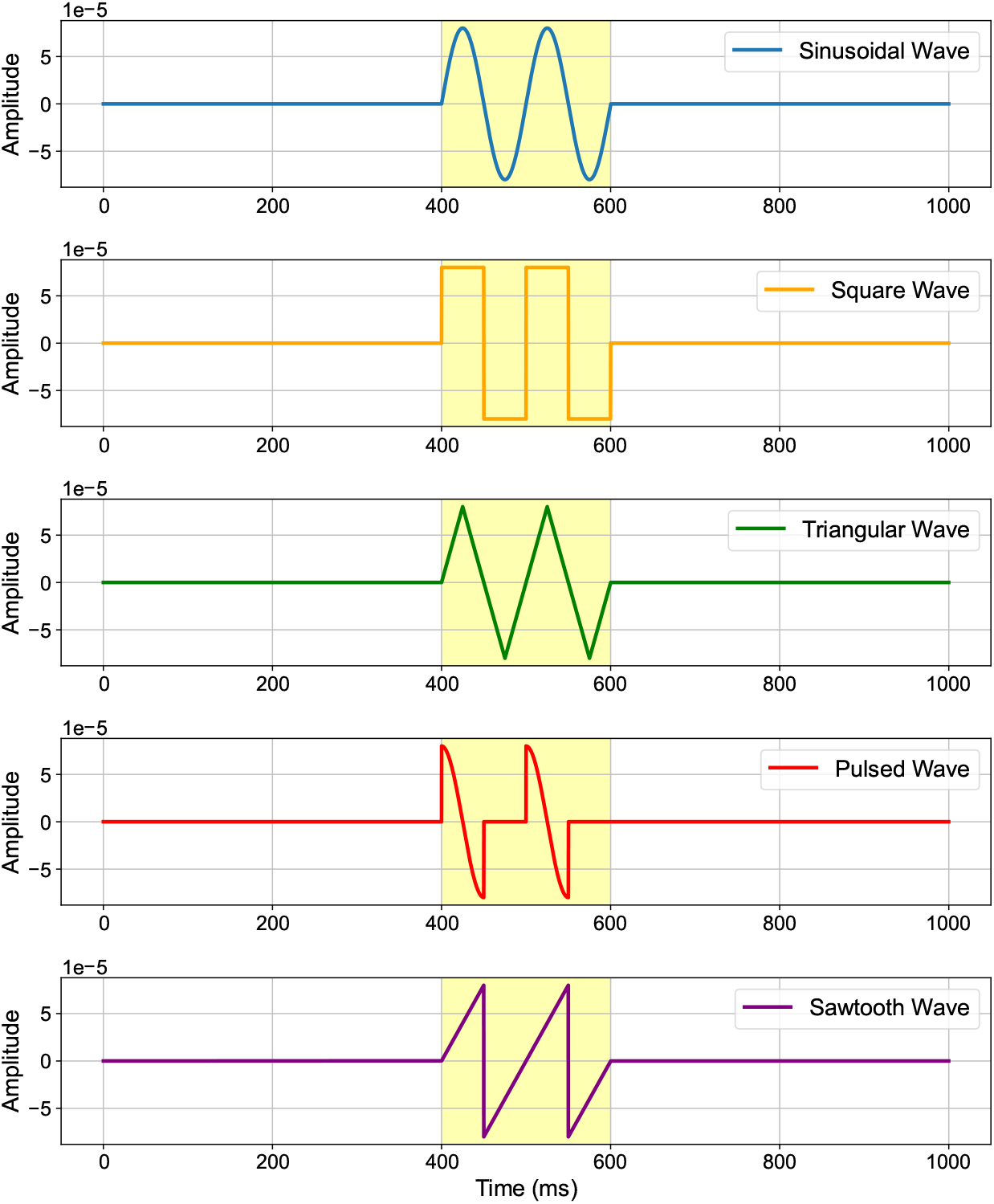
Representative perturbation waveforms used in the whole-brain simulations. Each panel shows the real part of the complex forcing term applied to the Stuart-Landau oscillators: (**a**) sinusoidal, (**b**) square, (**c**) triangular, (**d**) pulsed (50% duty cycle), and (**e**) sawtooth. For visualisation, all waveforms were plotted at the same peak amplitude and at *f* = 10 Hz, allowing direct comparison of their temporal profiles while preserving differences in their overall variance and harmonic content. The yellow shaded region highlights the 200 ms perturbation, including two cycles with a 100 ms period (10 Hz) interval shown for illustration, during which the external input was applied.

**Figure 2.**
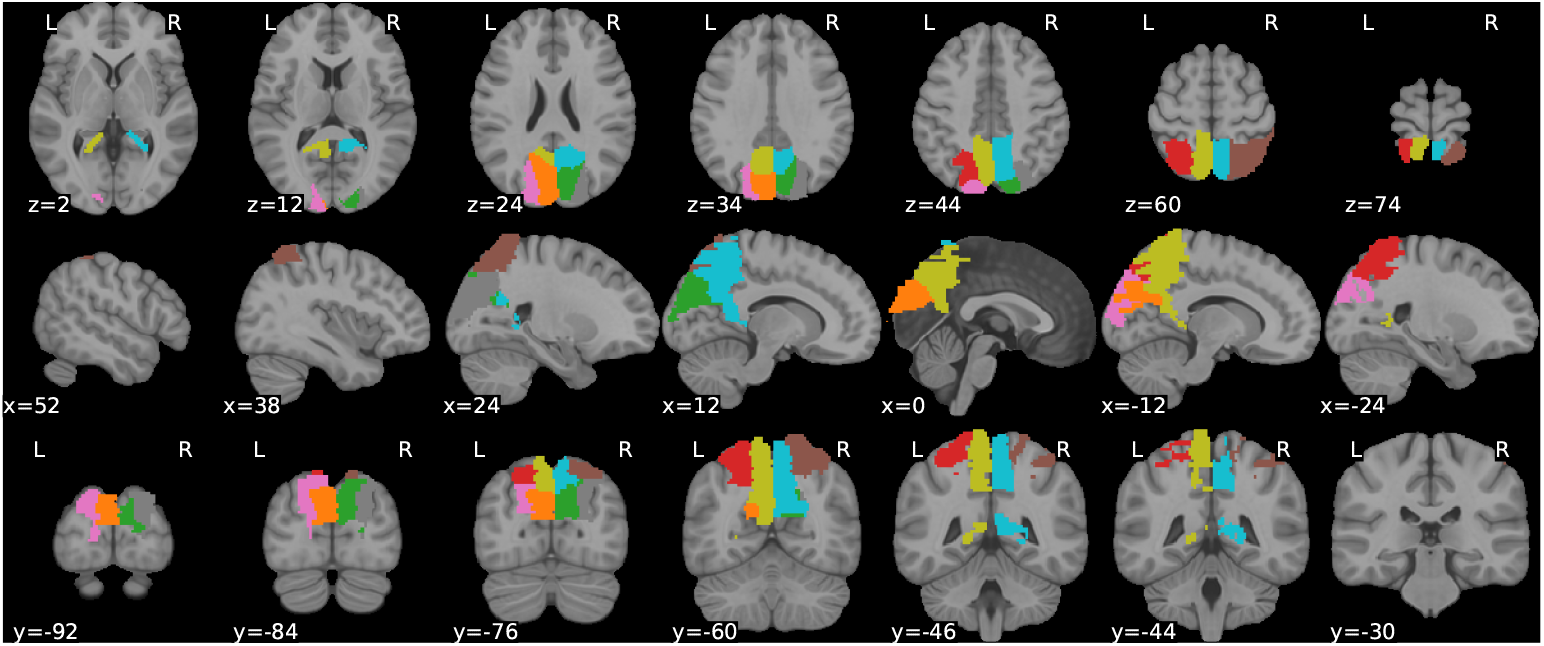
Anatomical MRI slices showing the posterior AAL regions selected for perturbation in the whole-brain model. Coloured overlays indicate the eight stimulated nodes: bilateral cuneus, superior occipital cortex, superior parietal cortex, and precuneus.

**Figure 3.**
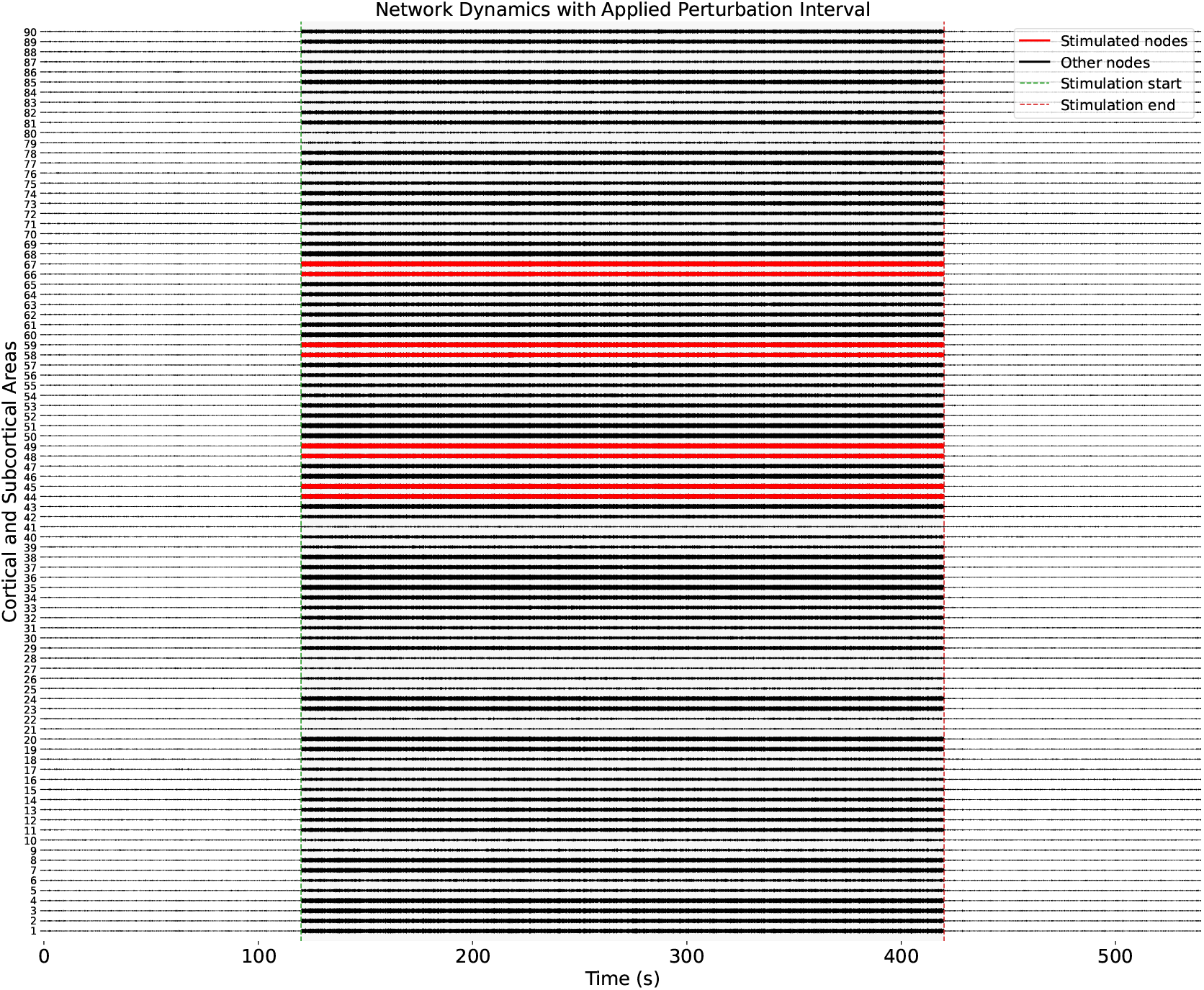
Hopf network dynamics with an externally applied perturbation interval. Time-series activity from all nodes is shown; the shaded region marks the perturbation window, and the stimulated nodes (44, 45, 48, 49, 58, 59, 66, 67) are highlighted in red during this period. A localised perturbation propagates through the Hopf network, modifying the collective dynamics relative to the unperturbed baseline.

- **Sinusoidal waveform**:

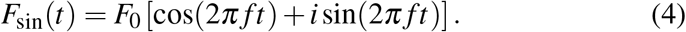
- **Square waveform**, implemented as a quadrature-shifted binary waveform:

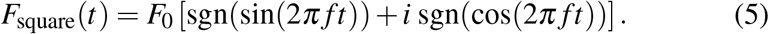

Here, sgn(*x*) = 1 for *x ≥* 0 and −1 otherwise.
- **Triangular waveform**, implemented as a symmetric triangular waveform with quadrature shift:

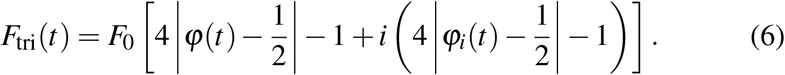
- **Pulsed waveform**, implemented as a gated sinusoidal drive with duty cycle *d* = 0.5:

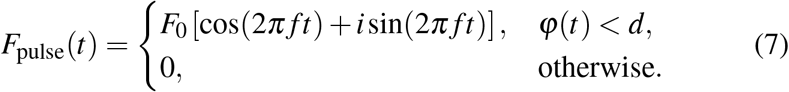
- **Sawtooth waveform**, implemented as a centred sawtooth with quadrature shift:

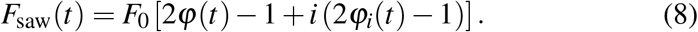

All waveforms were applied at a fixed stimulation frequency of *f*_stim_ = 11 Hz, which corresponded to the dominant spontaneous alpha peak of the fitted whole-brain model. This frequency was selected to target the model’s endogenous alpha resonance and was held constant across all waveform conditions so that observed differences in network dynamics could be attributed solely to waveform shape.

The stimulation amplitude was fixed at *F*_0_ = 2.1 × 10^−4^. This value was determined following the perturbational framework of [13], in which the strength of the external forcing is systematically varied to identify amplitudes that produce measurable changes in network dynamics while preserving the intrinsic operating regime of the model. In preliminary simulations, the forcing amplitude was therefore varied and the model response was quantified using alpha-band power enhancement and phase locking to the external drive. The selected amplitude produced a robust increase in alpha-band activity while maintaining the characteristic metastable dynamics of the unperturbed system.

Figure 1 illustrates the temporal profiles of the five forcing functions. Together, they span both smooth and discontinuous forms of rhythmic input, enabling a systematic assessment of how waveform geometry shapes entrainment and propagation in large-scale brain networks.

#### Propagation through the network

Once introduced, perturbations propagate across the system via the structural coupling and conduction delays defined by the connectome. The dynamics of entrainment are therefore determined not only by the local properties of the driven nodes, but also by the network topology and the timing of interactions. This framework allows us to systematically compare how different waveform shapes entrain oscillatory activity, amplify or suppress network synchrony, and modulate large-scale emergent dynamics.

### 1.5 Dynamical characterisation of simulated brain activity

All analyses were performed on the simulated time series from each of the 90 AAL brain regions after discarding an initial 60 s transient and standardising each regional time series (z-score). For each stimulation condition and waveform, two non-overlapping 120 s windows were analysed: *Rest* (0–120 s) and *Perturbed* (180–300 s), defined relative to the post-transient time series. Within each analysis window, signals were segmented into 5 s non-overlapping epochs (5,000 samples). Synchrony, metastability, and spectral entropy were computed per epoch and then summarised by their mean within each window. To guard against undue influence of rare extreme values, outlier rejection was applied to the per-epoch distributions using the interquartile range rule (threshold 1.5 × IQR). For statistical comparison between *Rest* and *Perturbed* conditions, paired Wilcoxon signed-rank tests with continuity correction were used when epoch counts matched; otherwise, two-sided Mann–Whitney *U* tests were applied.

#### Synchrony and Metastability

Large-scale phase coordination in the *α* band was quantified via the Kuramoto order parameter. Node signals were band-pass filtered (4th-order zero-phase Butterworth, 8–13 Hz), analytic phases obtained by the Hilbert transform, and unit phasors formed as 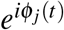 . The instantaneous global synchrony is

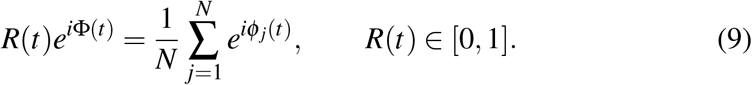

Two summary metrics were extracted per epoch: *synchrony S* = ⟨*R*(*t*) ⟩ _*t*_, the temporal mean of *R*(*t*), and *metastability M* = std_*t*_ *R*(*t*), the temporal dispersion of *R*(*t*) that captures the variability of network coordination [9, 12, 20]. These measures were computed on non-overlapping 5 s epochs and aggregated within *Rest*/*Perturbed* windows.

#### Spectral entropy (SE)

Spectral concentration within the *α* band was assessed using Shannon spectral entropy [21, 22]. For each 5 s epoch, the power spectral density was estimated using Welch’s method, and only frequencies within the alpha range (8–13 Hz) were retained. The band-limited spectrum *P*( *f*_*k*_) was normalized such that

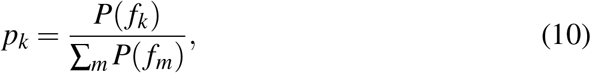

and the spectral entropy was computed as

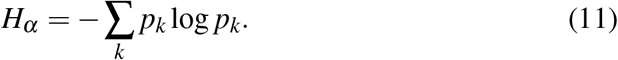

This quantity was calculated separately for each of the 90 AAL regions and then averaged across regions to obtain a single value per epoch. Lower *H*_*α*_ indicates that power is concentrated around a narrow set of frequencies (a more peaked alpha spectrum), whereas higher *H*_*α*_ reflects a broader and flatter spectral distribution.

#### Metastable oscillatory modes (MOMs)

To quantify transient, band-limited episodes of elevated oscillatory activity, we analysed metastable oscillatory modes (MOMs). We followed a threshold-based envelope methodology grounded in whole-brain modelling work on delay-coupled oscillators [23]. For each node *n*, band-pass filtering (4th-order zero-phase Butterworth, 8–13 Hz) was applied to the real component of the simulated signals, and the analytic signal was obtained via the Hilbert transform. The instantaneous *α*-band amplitude (envelope) is then *A*_*n*_(*t*) = |*ℋ* {*x*_*n*_(*t*)} |.

To avoid circularity, thresholds were estimated exclusively from the *Rest* window and then held fixed when scoring both *Rest* and *Perturbed* windows. Specifically, per-node thresholds were defined as 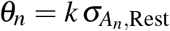, where 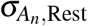 is the standard deviation of the baseline alpha-band envelope. A single multiplicative factor *k* was selected so that the resulting thresholds yielded a global baseline occupancy in the target range of 1–15%. As a robustness check, a percentile rule (*θ*_*n*_ equal to the 95th percentile of *A*_*n*_ in *Rest*) produced qualitatively similar results. Binary MOM masks were defined as

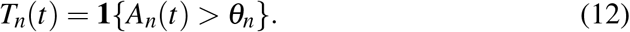

A minimum duration of two *α* cycles was enforced, removing suprathreshold runs shorter than ≈ 0.2–0.25 s to exclude spurious, ultrashort excursions.

From {*T*_*n*_(*t*)} we derived three standard MOM descriptors per condition: *occupancy, burst count*, and *burst duration*. Occupancy was computed per node as the fraction of time above threshold,

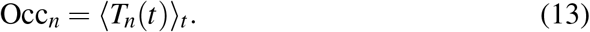

Occupancy values were then summarised across nodes. Burst counts were obtained by enumerating contiguous suprathreshold runs in *T*_*n*_(*t*) per node. Burst durations (in seconds) were computed as run lengths divided by the sampling rate and then pooled across nodes to characterise the distribution of episode lengths. This operationalisation mirrors prior work in which MOMs are defined as transient, spatially extended increases in band-limited amplitude emerging from weakly stable cluster synchronisation in delay-coupled networks, and whose occupancy, size and duration are systematically controlled by global coupling and conduction delays [23].

## 2 Results

### Metastability and synchrony under temporally structured perturbations

We quantified how waveform shape alters macroscopic network dynamics in the Stuart–Landau whole-brain model operating on the subcritical side of the Hopf bifurcation. For each perturbation, metastability, mean synchrony, and spectral entropy were computed in the *Rest* and *Perturbed* windows (Fig. 4).

**Figure 4:**
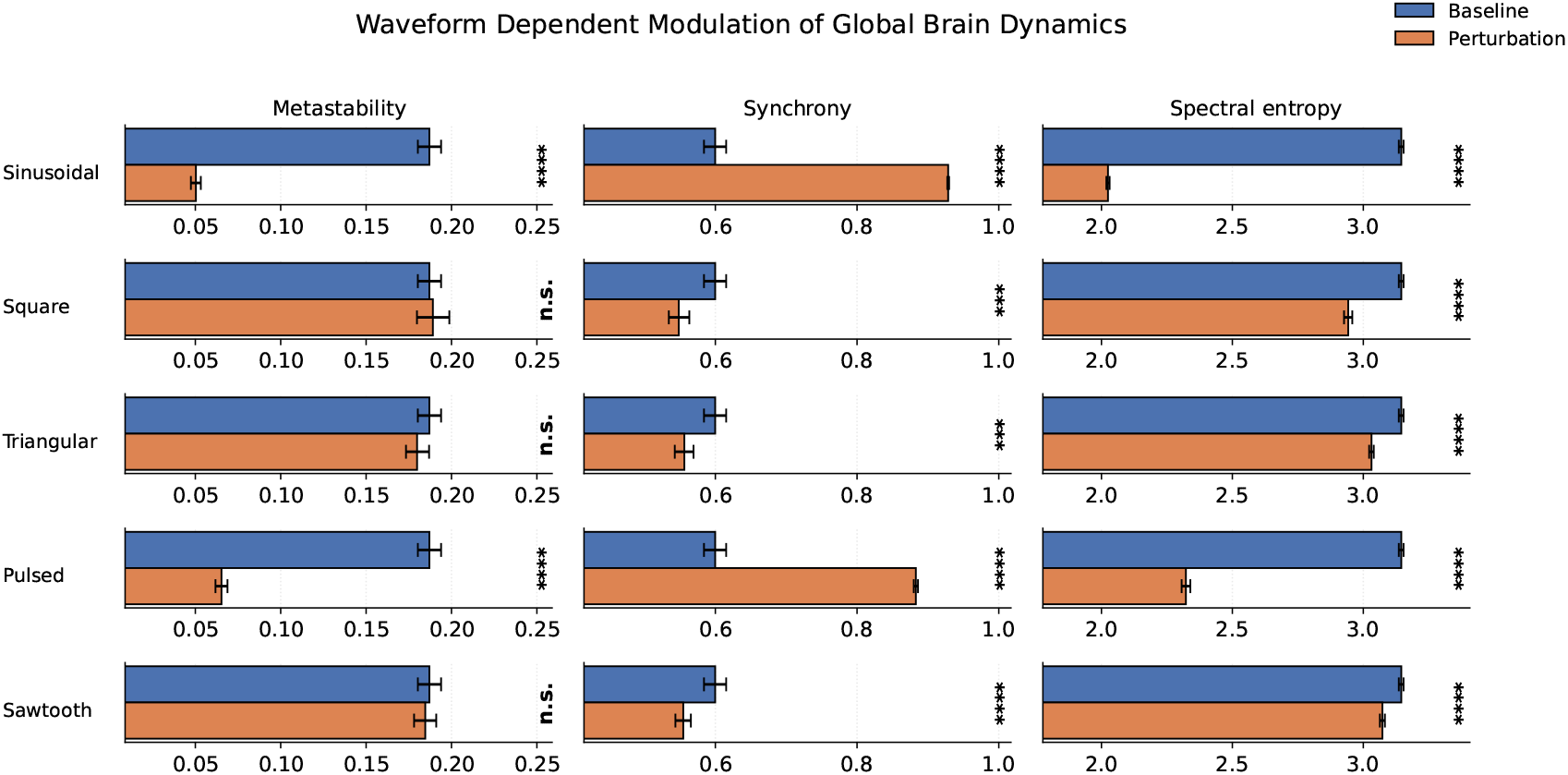
Waveform shape determines global synchronisation in the whole-brain model. Metastability (*σ* (KOP)), synchrony ( ⟨ KOP ⟩), and spectral entropy are shown for the *Rest* (blue) and *Perturbed* (orange) windows for five perturbation waveforms. Sinusoidal and pulsed stimulation produce large reductions in metastability and spectral entropy, accompanied by pronounced increases in synchrony, indicating a transition toward a highly coherent and low-complexity dynamical state. In contrast, square, triangular, and sawtooth waveforms do not significantly alter metastability, but reduce synchrony and produce more modest decreases in spectral entropy. Bars show means with bootstrap 95% confidence intervals; asterisks denote statistical comparisons between Rest and Perturbed periods.

#### 2.0.1 Sinusoidal and pulsed stimulation drive robust global state transitions

A clear separation emerged between waveform classes. Sinusoidal and pulsed perturbations produced the strongest and most consistent changes across all metrics. During sinusoidal stimulation, metastability decreased from 0.187 to 0.050 (*p <* 10^−6^), while mean synchrony increased from 0.600 to 0.929 (*p <* 10^−6^). Pulsed stimulation produced a similarly pronounced effect, reducing metastability to 0.065 and increasing synchrony to 0.883 (both *p <* 10^−6^). Spectral entropy decreased substantially under both waveforms, falling from 3.15 to 2.02 for sinusoidal stimulation and to 2.32 for pulsed stimulation (*p <* 10^−8^). Together, these changes indicate a strong transition from a fluctuation-rich metastable regime toward a highly coherent and spectrally constrained network state.

#### 2.0.2 Square, triangular, and sawtooth waveforms preserve metastability but reduce synchrony

In contrast, square, triangular, and sawtooth perturbations did not produce significant changes in metastability. Square stimulation left metastability essentially unchanged (0.187 to 0.189, *p* = 0.90), triangular stimulation produced a small non-significant decrease (0.187 to 0.178, *p* = 0.060), and sawtooth stimulation similarly showed no reliable effect (0.187 to 0.185, *p* = 0.66). Despite preserving metastability, all three waveforms significantly reduced mean synchrony, decreas-ing it from approximately 0.600 at baseline to 0.544–0.556 during stimulation (all *p <* 0.001). Spectral entropy also decreased significantly, although the magnitude of these reductions was modest compared with those observed for sinusoidal and pulsed stimulation. These results indicate that waveforms with sharper transitions alter the spectral organisation of the system without collapsing its intrinsic temporal variability.

#### 2.0.3 Waveform shape determines the magnitude and direction of macroscopic state transitions

Taken together, these findings demonstrate that waveform geometry critically determines how rhythmic perturbations reshape large-scale brain dynamics. Sinusoidal and pulsed stimulation drive the system toward a strongly synchronised, low-entropy attractor characterised by markedly reduced metastability. In contrast, square, triangular, and sawtooth waveforms largely preserve the metastable operating regime while reducing global phase coherence.

##### Dynamical regime transition

The concurrent reductions in metastability and spectral entropy observed under sinusoidal and pulsed stimulation indicate a substantial narrowing of the system’s dynamical repertoire, consistent with a shift away from a fluctuation-rich regime toward a more ordered state. These changes suggest that temporally structured forcing can stabilise global oscillations by suppressing amplitude and phase variability, in line with theoretical predictions for phase-coherent perturbations near the Hopf bifurcation.

Across waveform conditions, changes in metastability and synchrony exhibited a strong inverse relationship, indicating that reductions in temporal variability were tightly coupled to increases in global phase alignment. Together, these findings support a mechanistic interpretation in which stimulation waveform shape determines whether the network is driven into a coherent oscillatory attractor or remains within a flexible metastable regime.

### 2.1 Stimulation waveform shapes metastable oscillatory modes

To characterise how perturbations reshape transient alpha activity, we quantified metastable oscillatory modes (MOMs) by measuring burst occupancy, burst count, and burst duration in the *Rest* and *Perturbed* windows (Fig. 5).

**Figure 5:**
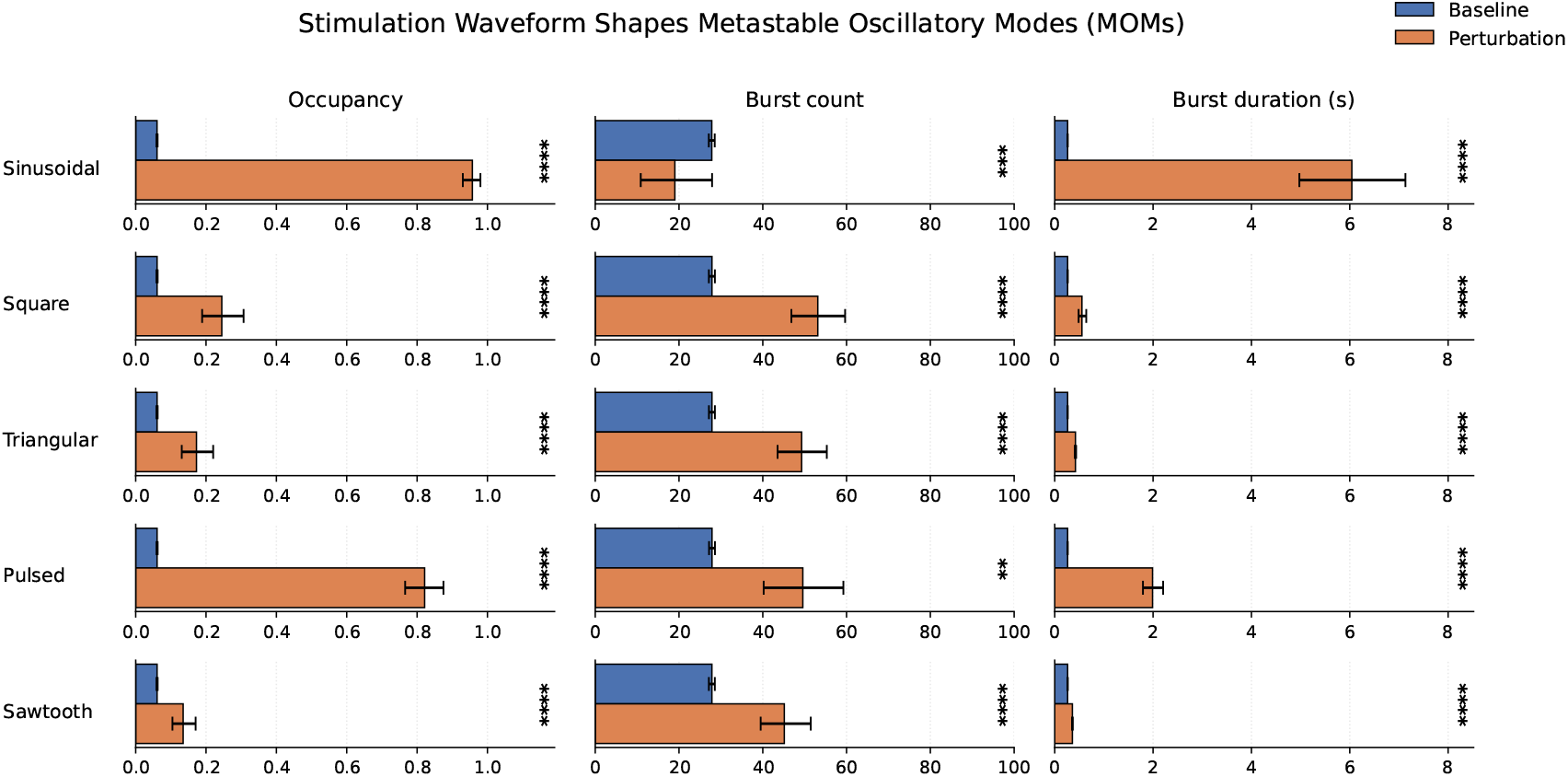
Waveform shape determines the transition from transient bursting to sustained oscillations. Burst occupancy, burst count, and burst duration are shown for the *Rest* (blue) and *Perturbed* (orange) windows for five perturbation waveforms. Sinusoidal stimulation drives near-continuous oscillatory activity, characterised by maximal occupancy, strongly prolonged burst duration, and a sharp reduction in burst count. Pulsed stimulation produces highly occupied and long-lasting bursts, but without a corresponding reduction in burst count, indicating a more fragmented entrainment pattern. Square, triangular, and sawtooth waveforms produce only modest increases in burst occupancy and duration while increasing burst count, consistent with preservation of the metastable bursting regime. Bars show means with bootstrap 95% confidence intervals; asterisks denote statistical comparisons between Rest and Perturbed periods.

#### 2.1.1 Sinusoidal stimulation induces near-continuous oscillatory activity

Sinusoidal stimulation produced the most pronounced change in bursting dynamics. Burst occupancy increased from 0.061 to 0.9999 (*p <* 10^−28^), indicating that oscillatory activity became nearly continuous throughout the perturbation period. In parallel, burst count decreased dramatically from 27.8 to 1.2 (*p <* 10^−28^), while burst duration increased from 0.246 s to 0.653 s (*p <* 0.001). Together, these changes indicate a transition from intermittent metastable bursts to a sustained oscillatory regime.

#### 2.1.2 Pulsed stimulation strongly prolongs bursts without reducing burst count

Pulsed stimulation produced a closely related pattern. Burst occupancy increased markedly from 0.061 to 0.884 (*p <* 10^−28^), and burst duration increased from 0.246 s to 0.636 s (*p <* 0.001). In contrast to sinusoidal stimulation, burst count did not change significantly (27.8 to 49.6, *p* = 0.62), reflecting substantial variability across epochs. These results indicate that pulsed stimulation promotes long-lasting oscillatory episodes while preserving a more fragmented temporal structure than the sinusoidal waveform.

#### 2.1.3 Square, triangular, and sawtooth waveforms produce modest changes in bursting

Square, triangular, and sawtooth waveforms produced significantly smaller changes. Burst occupancy increased only modestly, reaching 0.132 for square, 0.090 for triangular, and 0.075 for sawtooth stimulation. Burst durations increased slightly from 0.246 s at baseline to approximately 0.284–0.296 s during stimulation. Burst count increased substantially for all three waveforms, rising to 47.9 (square), 39.0 (triangular), and 34.8 (sawtooth), indicating more frequent but still short-lived oscillatory episodes. Thus, these waveforms enhanced burst occurrence without inducing sustained oscillatory activity.

#### 2.1.4 Waveform shape determines the transition from metastable bursting to sustained oscillations

These results mirror the Kuramoto order parameter findings and demonstrate that waveform shape critically determines how stimulation reshapes transient oscillatory dynamics. Sinusoidal stimulation drove the system into a near-continuous oscillatory state, while pulsed stimulation produced prolonged and highly occupied bursts with greater temporal fragmentation. In contrast, square, triangular, and sawtooth waveforms preserved the metastable bursting regime, producing only modest increases in occupancy and duration despite more frequent burst events.

## 3 Discussion

In this work, we used a whole-brain network of delay-coupled Stuart–Landau oscillators to examine how the *shape* of externally applied rhythmic stimulation modulates large-scale brain dynamics. By systematically comparing sinusoidal, square, triangular, pulsed, and sawtooth perturbations at matched frequency and amplitude, we show that waveform geometry is a key determinant of macroscopic network behaviour. Across multiple complementary metrics, including global synchrony, metastability, spectral entropy, and the statistics of transient alpha-band bursts, we find that only sinusoidal and pulsed stimulation reliably drive the system into a highly synchronised, low-entropy regime, whereas square, triangular, and sawtooth waveforms produce weaker and qualitatively different effects despite being delivered at the same frequency and amplitude.

### Waveform-dependent entrainment and collective dynamics

A central finding of this study is that sinusoidal and pulsed perturbations robustly increased global synchrony while concomitantly reducing metastability and spectral entropy. From a dynamical-systems perspective, this pattern reflects a shift from itinerant dynamics near the edge of criticality toward a more ordered regime dominated by coherent phase alignment [9, 10]. Operating near the Hopf bifurcation renders each node highly susceptible to external forcing [10]; however, the *temporal profile* of the drive determines how efficiently phase relationships are recruited and stabilised across the network.

Although pulsed and sawtooth waveforms both contain sharp temporal transitions and enriched harmonic structure [6, 7], they produced markedly different network responses. Pulsed stimulation induced strong entrainment comparable to sinusoidal forcing, with pronounced reductions in metastability and spectral entropy and a substantial increase in global synchrony. In contrast, sawtooth stimu-lation did not significantly alter metastability and instead produced a reduction in synchrony together with only a modest decrease in spectral entropy. This dissociation indicates that harmonic richness alone is insufficient to predict large-scale network effects; rather, the precise temporal arrangement of the waveform critically determines how effectively phase relationships are stabilised across the network.

Square, triangular, and sawtooth waveforms produced weaker and more fragmented modulation of global coordination. Although these waveforms also contain higher harmonics, their temporal structure was less effective at stabilising coherent phase relationships in the present network configuration. Instead, they reduced global synchrony and produced only partial reductions in spectral entropy, while metastability remained largely unchanged. These results underscore that wave-form shape influences not merely the presence of harmonics but also how phase information is temporally structured and integrated by the network.

### Metastability, criticality, and dynamical regime transitions

The concurrent decrease in metastability and increase in synchrony under sinusoidal and pulsed stimulation indicate a reduction in the temporal variability of global coordination. In prior whole-brain modelling work, high metastability has been associated with flexible switching between partially synchronised states, a hallmark of dynamics poised near criticality [9, 10, 12]. In this context, our results suggest that appropriately structured stimulation can transiently displace the system away from this regime, stabilising a smaller subset of attractors characterised by strong coherence.

The strong negative relationship between changes in metastability and synchrony across waveform conditions supports this interpretation. Rather than acting as independent dimensions, these metrics jointly reflect how external forcing reshapes the stability landscape of the network. Importantly, this transition is highly waveform-specific and arises within the physiological operating regime of the model, highlighting how stimulation can be used as a controlled probe of network susceptibility. This aligns with perturbational perspectives on brain dynamics, in which the system’s response to weak external input reveals latent structure in its attractor landscape [13].

### Alpha bursts as signatures of waveform-driven regime changes

The analysis of metastable oscillatory modes provides a complementary view of these effects at the level of transient amplitude dynamics. Under baseline conditions, alpha activity manifested as sparse, short-lived bursts distributed across nodes, consistent with prior modelling and empirical work linking metastable os-cillatory modes to weakly stable cluster synchronisation [23]. Effective waveforms dramatically altered this pattern.

Sinusoidal stimulation strongly increased alpha power, driving alpha envelopes continuously above baseline threshold. This resulted in occupancy values approaching unity while reducing burst count as discrete episodes merged into sustained activity. Pulsed stimulation produced similarly large increases in occupancy and burst duration, but retained higher burst counts, indicating repeated and prolonged episodes rather than complete saturation. In contrast, square, triangular, and saw-tooth waveforms primarily increased burst count while producing comparatively small changes in occupancy and duration, consistent with more fragmented and transient oscillatory activity.

These distinctions mirror the global synchrony and entropy findings, reinforcing the interpretation that waveform shape controls not only phase coherence but also the temporal organisation of amplitude fluctuations. In this sense, metastable oscillatory mode statistics provide an intuitive bridge between local oscillatory phenomena and global coordination measures, linking transient bursts to shifts in large-scale dynamical regimes. These findings suggest that waveform-dependent changes in global synchrony emerge from the reorganisation of local burst dynamics rather than from phase alignment alone.

### Relation to empirical stimulation studies

Although the present results are derived from a computational model, they align with and extend existing empirical findings on waveform-specific neuromodulation. Experimental studies have shown that non-sinusoidal stimulation, particularly pulsed and other temporally structured variants, can alter excitability and entrainment beyond what is achieved with conventional sinusoidal stimulation alone [6–8]. Our results provide a mechanistic account for these observations, suggesting that waveform-dependent phase-response interactions at the local level scale up to network-wide reconfiguration when embedded in realistic structural connectivity.

Importantly, the model isolates neural dynamics from stimulation artefacts, allowing for a clean comparison across waveform designs. The strong and consistent effects observed here therefore constitute testable predictions for human EEG and MEG studies, particularly regarding changes in metastability, spectral entropy, and burst statistics under different stimulation waveforms.

### Limitations and future directions

Several limitations should be acknowledged. First, the model employs a homogeneous local oscillator architecture and does not explicitly represent laminar structure or cell-type-specific responses, which may modulate waveform sensitivity in biological tissue. Second, stimulation was applied to a fixed set of posterior visual–parietal nodes at a single frequency and amplitude; future work should explore how waveform effects interact with spatial targeting, frequency detuning, and state dependence. Finally, while the present study focuses on alpha-band dynamics, non-sinusoidal waveforms may differentially affect cross-frequency interactions, an avenue that remains unexplored here.

Future extensions could incorporate adaptive or closed-loop waveform design [24], leveraging real-time estimates of synchrony or metastability to steer the system toward desired dynamical states. More broadly, combining whole-brain modelling with empirical stimulation data will be essential for validating and refining the mechanistic predictions advanced here.

### Implications

Taken together, our findings demonstrate that waveform shape is not a secondary implementation detail but a fundamental determinant of how rhythmic stimulation interacts with large-scale brain dynamics. By shaping phase-response interactions and attractor stability, stimulation waveforms can selectively promote either sustained network entrainment, as observed for sinusoidal and pulsed stimulation, or more fragmented and weakly coordinated dynamics, as observed for square, triangular, and sawtooth stimulation. These results position waveform design as a principled lever for network-level neuromodulation, extending the traditional focus on frequency and amplitude and opening new avenues for both basic neuroscience and clinical intervention.

## Acknowledgements

VS and PT acknowledge support from the European Union through AISN (Horizon Europe, Grant No. 101057655) and eBRAINS-Health (Horizon Europe, Grant No. 101058516). JC was supported by LARSyS funding from the Portuguese Foundation for Science and Technology (FCT, Portugal; DOI: 10.54499/LA/P/0083/2020).

## Data availability

The data generated and analysed during the current study are available upon reasonable request to Vivek Sharma.

## Code availability

The code used for simulations and analysis in this study is currently maintained in a private GitHub repository for the purpose of peer review and will be made publicly available upon publication.

## Author information

### Contributions

J.C., P.H.E.T., and V.S. conceived the study. V.S. implemented the simulations and performed the analyses. J.C. contributed to the modelling framework and analysis scripts. V.S. and J.C. developed the modelling framework and interpreted the results. P.H.E.T. provided theoretical guidance and supervised the project. V.S. wrote the original manuscript draft. All authors reviewed, edited, and approved the final version of the manuscript.

